# Insulin-like growth factor receptor / mTOR signaling elevates global translation to accelerate zebrafish fin regenerative outgrowth

**DOI:** 10.1101/2022.10.09.511495

**Authors:** Victor M. Lewis, Heather K. Le Bleu, Astra L. Henner, Hannah Markovic, Amy E. Robbins, Scott Stewart, Kryn Stankunas

## Abstract

Zebrafish robustly regenerate fins, including their characteristic bony ray skeleton. Amputation activates intra-ray fibroblasts and dedifferentiates osteoblasts that migrate under a wound epidermis to establish an organized blastema. Coordinated proliferation and re-differentiation across lineages then sustains progressive outgrowth. We generate a single cell transcriptome dataset to characterize regenerative outgrowth and explore coordinated cell behaviors. We computationally identify sub-classes of fibroblast-lineage cells and describe novel markers of intra- and inter-ray fibroblasts and growth-promoting distal blastema cells. A pseudotemporal trajectory and in vivo photoconvertible lineage tracing indicate distal blastemal mesenchyme generates both re-differentiated intra- and inter-ray fibroblasts. Gene expression profiles across this trajectory suggest elevated protein translation in the blastemal mesenchyme state. O-propargyl-puromycin incorporation and small molecule inhibition identify insulin growth factor receptor (IGFR) / mechanistic target of rapamycin kinase (mTOR)-dependent elevated bulk translation in blastemal mesenchyme and differentiating osteoblasts. We test candidate cooperating differentiation factors identified from the osteoblast trajectory, finding IGFR/mTOR signaling expedites glucocorticoid-promoted osteoblast differentiation *in vitro*. Concordantly, mTOR inhibition slows but does not prevent fin regenerative outgrowth *in vivo*. IGFR/mTOR may elevate translation in both fibroblast- and osteoblast-lineage cells during the outgrowth phase as a tempo-coordinating rheostat.

## INTRODUCTION

Unlike mammals, most teleost fish robustly regenerate appendages following injury. The zebrafish caudal fin provides a tractable system to investigate fin regeneration mechanisms. Injured caudal fins return to their original form in approximately 3 weeks, recapitulating uninjured size, shape and tissue organization. Fins are scaffolded by a bony ray (or lepidotrichia) skeleton. Fibroblasts, vasculature, and sensory axons reside within and between rays, A stratified epidermis surrounds both rays and inter-ray tissue. Signaling coordinates behaviors of the varied fin cell types, including fibroblasts and bone-producing osteoblasts, to initiate regeneration and progressively restore rays and other fin tissue (reviewed in Sehring and Weidinger, 2019).

Injury “activates” intra-ray fibroblasts that migrate distally and proliferate to contribute the core blastemal mesenchyme during the establishment phase of fin regeneration. The distal blastemal mesenchyme pool transitions into a morphologically distinct and growth factor-producing organizing center, or “niche”, to launch the outgrowth phase between two and three days post amputation (Nechiporuk & Keating, 2002, Wehner et al., 2014, Stewart et al., 2014, Mateus, et al., 2015, Tornini et al. 2016, Stewart et al. 2019). Throughout outgrowth, blastemal mesenchymal cells proliferate and differentiate to restore mature fibroblasts. Therefore, fibroblast-lineage cells are crucial to initiating a regenerative response, coordinating balanced outgrowth, and replacing lost connective tissue.

Epithelialized osteoblasts proximal to the wound site dedifferentiate to a mesenchymal pre-osteoblast (pOb) state and, in parallel with fibroblasts, contribute to the blastema (Knopf et al., 2011, Tu and Johnson, 2011, Sousa et al., 2011, Stewart and Stankunas, 2012, Singh et al. 2012, Stewart et al., 2014). Compartmentalization within the blastema spatially restricts opposing pOb proliferation and differentiation activities while positioning pObs to restore ray pattern (Nechiporuk and Keating, 2002, Brown et al., 2009, Stewart et al., 2014, Armstrong et al., 2017). Dedifferentiated pObs line the blastema laterally and are maintained distally by Wnt/β-catenin signaling (Brown et al. 2009, Stewart et al., 2014, Wehner et al., 2014). Proximal, differentiating osteoblasts produce bone morphogenetic proteins (BMPs) that promote re-differentiation, at least in part by opposing Wnt activity (Smith et al. 2006, Stewart et al. 2014). However, the specific mechanisms by which these and other signals regulate osteoblast state transitions during fin regenerative outgrowth are incompletely resolved. Likewise, signals that coordinate the behaviors of osteoblast- and fibroblast-lineage cells for progressive and tempo-modulated regeneration of fin rays are unknown.

We produce a single cell transcriptomic dataset to characterize cell types, states and transition-driving mechanisms during fin regenerative outgrowth. We identify novel markers of fibroblast-lineage cells including distinguishing factors of intra- and inter-ray fibroblasts. Pseudotemporal trajectories and photoconvertible lineage tracing suggest inter-ray fibroblasts derive from distal-lateral blastemal mesenchyme cells. Pseudotime gene expression analyses led us to uncover greatly elevated insulin growth factor receptor (IGFR) and mechanistic target of rapamycin kinase (mTOR)-dependent bulk protein translation in blastemal mesenchyme and differentiating osteoblasts. In vitro, IGFR/mTOR accelerates glucocorticoid-promoted fin osteoblast differentiation. In vivo, mTOR activity appears to coordinate output of fibroblast- and osteoblast-lineage blastemal cells to modulate regenerative tempo. As such, IGFR/mTOR signaling represents a plausible environment-sensing and/or systemic influence on fin regenerative outgrowth.

## RESULTS

### Single cell transcriptomic cluster analysis identifies osteoblasts and two fibroblast-lineage subclasses

We evaluated the transcriptomes of 7224 individual cells collected from regenerating fins at the onset of the outgrowth phase (3 days post amputation [dpa]; 2756 cells) and during progressive outgrowth (7 dpa; 4468 cells) (Figure 1A). We generated a combined dataset comprising eleven clusters following dimensionality reduction and unsupervised clustering using Monocle3 (Cao et al., 2019) (Figure Supplement 1-1A-F). Established markers identified the majority of lineages contributing to regenerating fins, including a combined osteoblast/fibroblast cluster (Figure Supplement 1-1G, Figure 1B) (Farnsworth, 2020, Hou et al., 2020, Lewis et al., 2019, Saunders et al., 2019, Stewart et al., 2014).

**Figure 1.**
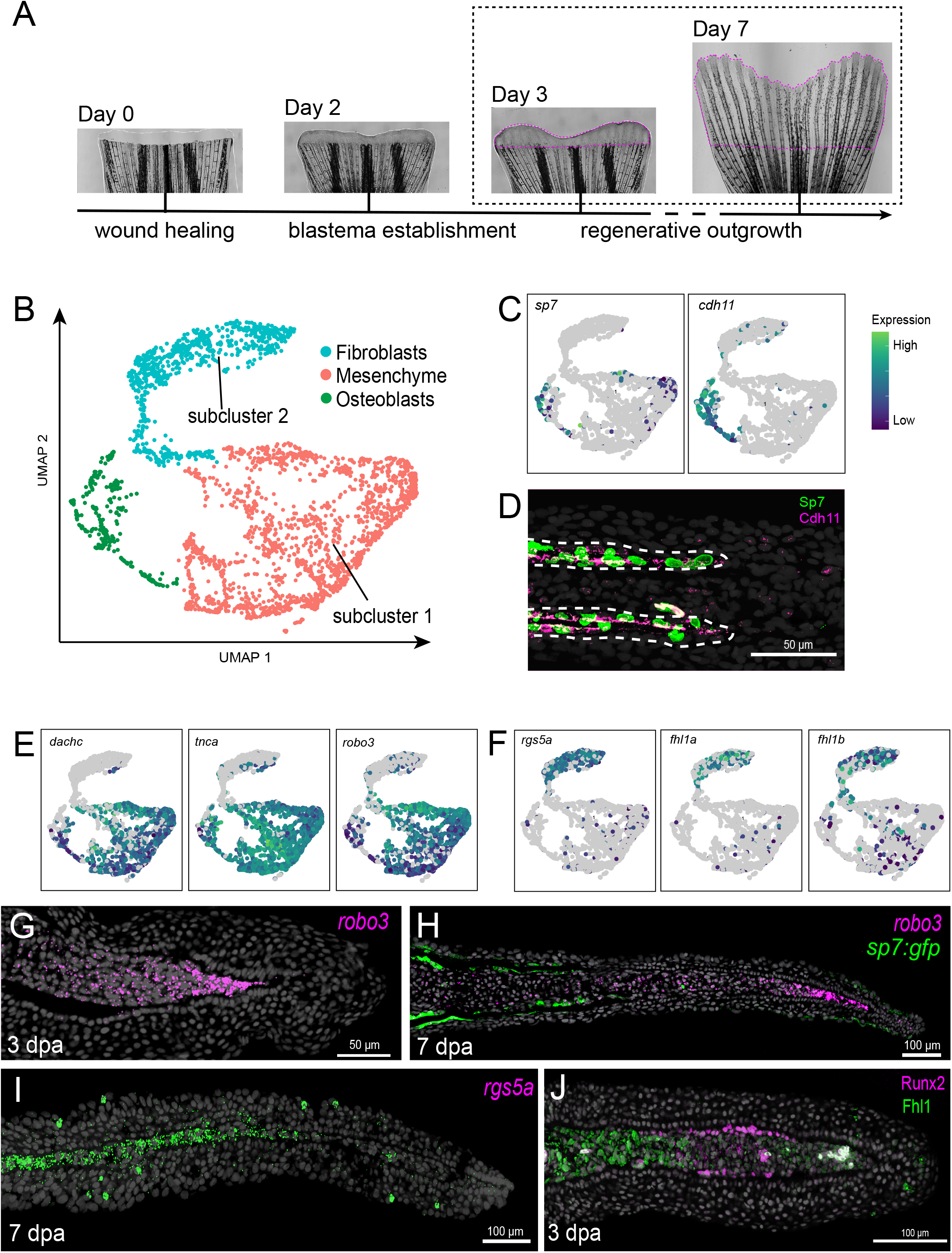
scRNA-Seq transcriptomics of zebrafish caudal fin regenerative outgrowth identifies distinguishing fibroblast and blastemal mesenchyme markers. (A) Whole mount images of regenerating fins highlighting collection of regenerating fin tissue from 3 and 7 days post amputation (dpa) for single-cell RNA sequencing (scRNAseq). Magenta dashed lines outline equivalent tissue used for single cell preparations. (B) UMAP plot of integrated datasets showing cell lineage and sub-class assignments of the fibroblast/osteoblast cluster. (C) UMAP plots showing co-localization of cells expressing the osteoblast markers *Sp7 transcription factor* (*sp7*) and *cadherin 11, type 2, OB-cadherin* (*cdh11*). (D) Antibody stained 3 dpa fin section showing co-localized Sp7 (green) and Cdh11 (magenta). (E) UMAP plots showing markers of: distal blastema *dachshund c* (*dachc*), proximal-medial mesenchyme *tenascin Ca* (*tnc*) and newly identified distal mesenchyme *roundabout, axon guidance receptor, homolog 3* (*robo3*). (F) UMAP plots showing markers of fibroblast lineage subcluster 2: *regulator of G protein signaling 5a* (*rgs5a*), *four and a half LIM domains 1a* (*fhl1a*), and *four and a half LIM domains 1b* (*fhl1b*). (G, H) RNAscope *in situ* hybridization of 3 and 7 dpa regenerate fin sections showing *robo3* expression (magenta) in distal blastema cells. *Tg(sp7:Egfp)* expression in (H, green) marks osteoblasts proximal to *robo3+* distal blastemal mesenchyme. (I) RNAscope *in situ* hybridization of a 7 dpa regenerate section demonstrating *rgs5a* expression (green) in proximal regenerated inter-ray fibroblasts and rare, undefined cells within the superficial epidermis. (J) Antibody staining of a 3 dpa regenerate sections showing Fhl1 expression (green) in proximal regenerated intra-ray fibroblasts. Section staining images (D, G-J) are confocal maximum intensity projections. Hoechst-stained nuclei are grey. Scale bars are 50 or 100 *μ*M, as indicated.

We isolated the osteoblast/fibroblast cluster for further analysis, aiming to resolve the two related cell lineages and their state transitions during regeneration. We identified osteoblasts co-clustering within UMAP space and expressing the osteoblast marker *cadherin 11 type 2 OB-cadherin (cdh11)* (Tang et al., 2021) (Figure 1B, C). We verified this assignment by antibody staining showing co-expression of Cdh11 and the osteoblast transcription factor Sp7 (Figure 1D). We removed the *cdh11*-defined osteoblasts from the osteoblast/fibroblast cluster and reiteratively sub-clustered the remaining cells to distinguish two sub-classes of fibroblast lineage cells (Figure 1B). Established markers indicated subcluster 1 was blastemal mesenchyme (e.g., *tnc* and the distal “niche” marker *dachc* (Govindan and Iovine, 2015, Stewart et al., 2019); Figure 1E, Figure Supplement 1-2A). Other blastemal markers (e.g., *wnt5a, msx3;* (Stoick-Cooper et al., 2007, Akimenko et al., 1995, Smith et al., 2006)) also were elevated in subcluster 1, albeit less distinguishing than expected (Figure Supplement 1-2B). The few *sp7*-expressing cells within subcluster 1 could reflect low, “leaky” *sp7* expression in certain fibroblast-lineage cell states or pre-osteoblasts that share a UMAP space-driving mesenchymal and/or progenitor state program with fibroblast-lineage blastemal mesenchyme.

We surveyed differentially expressed genes (DEGs) between the fibroblast lineage sub-classes to identify additional distinguishing markers and to characterize subcluster 2 (Supplemental File 1). We then used RNAscope for select transcripts to align UMAP space with *in situ* expression patterns. Subcluster 1-enriched *roundabout axon guidance receptor homolog 3* (*robo3*) expression was upregulated in distal blastema at both 3 and 7 dpa, similar to the *dachc* distal marker (Figure 1E, G, H) (Stewart et al., 2019). Subcluster 2-enriched *regulator of G protein signaling 5a* (*rgs5a*) marked proximal inter-ray fibroblasts at 7 dpa, which are no yet re-formed at 3 dpa (Figure 1F, I; Figure Supplement 1-2C, D). *four and a half LIM domains 1a/b* (*fhl1a*.*b)* was upregulated within subcluster 2 fibroblasts (Figure 1F). Correspondingly, Fhl1 protein was strongly enriched in presumptive re-differentiating fibroblasts of the proximal blastema at 3 dpa (Figure 1J). Subcluster 2-expressed *slit homolog 2* (*slit2*) was up-regulated in proximal intra-ray fibroblasts and expressed in the basal epidermis (Figure Supplement 1-2B, D). Subcluster 2 cells were mostly from 7 dpa samples, consistent with relatively more re-differentiated fibroblasts at a later outgrowth phase (Figure Supplement 1-1H) (Hou *et al*., 2020). Collectively, we conclude fibroblast subcluster 1 represents blastemal mesenchyme and subcluster 2 comprises mixed intra- and inter-ray fibroblasts.

### Pseudotemporal analysis identifies transcriptional routes through mesenchyme/fibroblast transitions

Differential expression of blastema markers across UMAP subcluster 1 space suggested blastemal mesenchyme heterogeneity was incompletely resolved by sub-clustering. Therefore, we used pseudotemporal analysis to identify transcriptional routes through mesenchyme/fibroblast subclusters to further distinguish transcriptional states and transitions (Figure 2A). After profiling genes differentially expressed across pseudotime (Supplemental File 2), we initiated a trajectory within actively cycling cells of subcluster 1 (e.g. *pcna, mki67, mcm3*; Figure 2B). The trajectory traversed distal blastema cells (the UMAP space with highest *dachc, robo3, wnt5a*, and *msx3* expressing cells) and passed into subcluster 2 fibroblasts (Gerdes et al., 1983, Kelman 1997, Valverde et al., 2018). This trajectory matches the lineage-traced differentiation of blastemal mesenchyme cells to intra-ray fibroblasts during fin outgrowth (Tu et al., 2011, Stewart et al., 2012, Tornini et. al., 2016).

**Figure 2.**
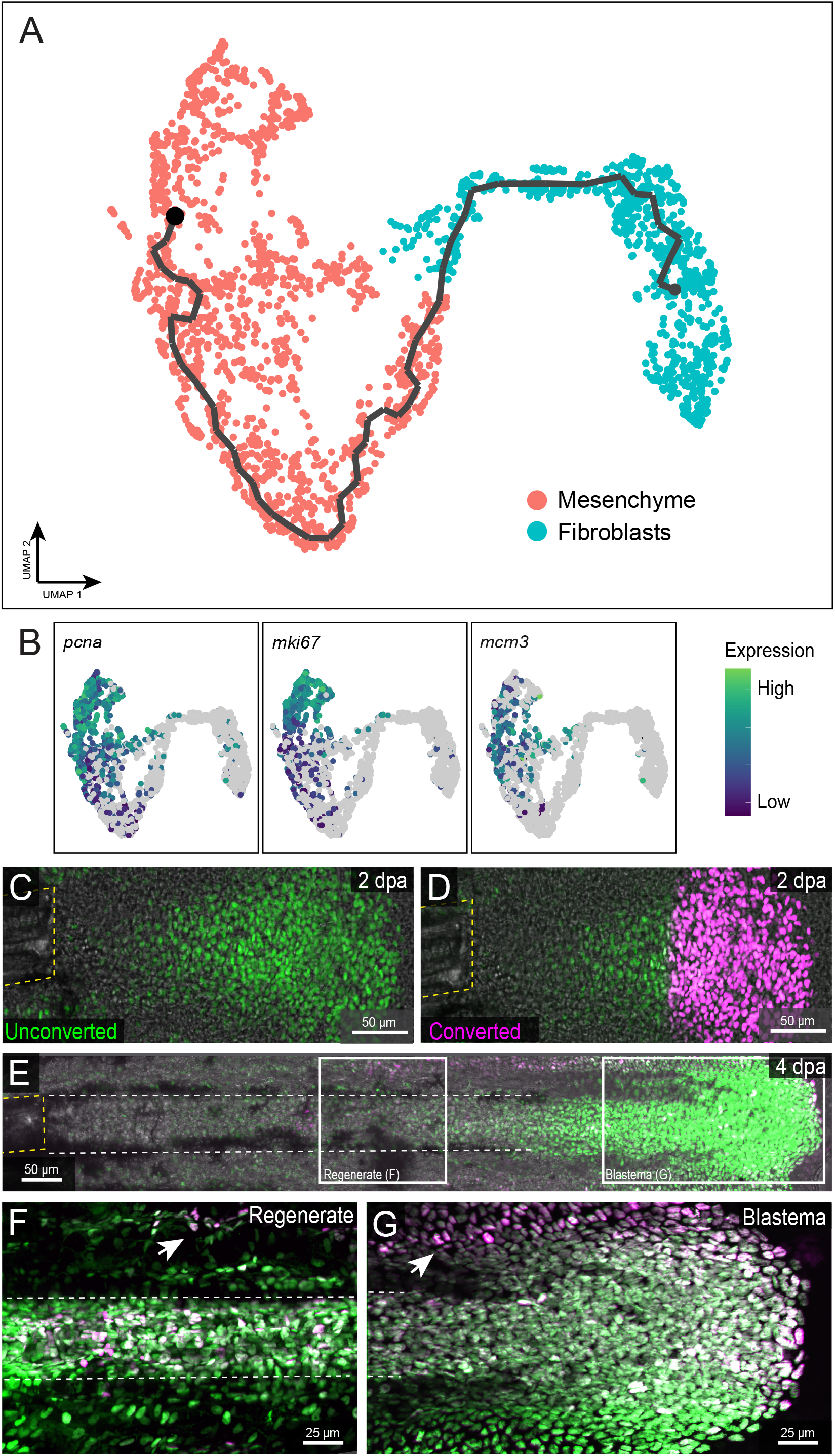
Intra- and inter-ray fibroblasts derive from distal blastemal mesenchyme. (A) UMAP plot with overlaid pseudotemporal trajectory traversing blastemal mesenchyme and fibroblasts. (B) UMAP plots showing relative expression of proliferation state markers *proliferating cell nuclear antigen* (*pcna*), *marker of proliferation Ki-67* (*mki67*) and *minichromosome maintenance complex component 3* (*mcm3*). The pseudotemporal trajectory in (A) initiates in proliferative mesenchyme cells. (C-G) Confocal whole mount caudal fin images of a *Tg(sost:nlsEos)* fish at 2 days post amputation (dpa) before (C) and after (D) Eos photoconversion and then re-imaged 48 hours later, at 4 dpa (E-G). Unconverted and photo- converted Eos is in green and magenta, respectively. (C, D) 20x images showing targeted Eos photo-conversion in distal blastemal mesenchyme cells following localized UV light exposure. (E) 20x stitched image showing proximalization of photo-converted Eos-expressing cells 48 hours post conversion (at 4 dpa). (F, G) 40x images of zoomed “regenerate” and “blastema” regions shown in (E). Photo-converted Eos-expressing inter- and intra-ray regenerated fibroblasts are present proximally (F) and appear to migrate from the blastema toward inter-ray regenerate regions (G). Yellow dashed lines outline the bony ray stump and mark the amputation plane. White dashed lines highlight regenerating rays at 4 dpa. Arrows mark inter-ray fibroblasts expressing photo- converted Eos. Scale bars are 25 or 50 *μ*M, as indicated.

The pseudotemporal trajectory suggests blastemal mesenchyme also produces inter-ray fibroblasts, as indicated by Cre/lox-based lineage tracing (Tornini et al., 2016). We further evaluated this lineage relationship using the Tg(*sost:nlsEos*) photoconvertible fluorescent line (Thomas and Raible, 2018), which we characterized as labeling non-osteoblast blastemal mesenchyme and regenerated intra- and inter-ray fibroblasts (Figure Supplement 2-1). We photoconverted blastema mesenchyme at 2 dpa (Figure 2C, D), prior to fibroblast re-differentiation, and observed converted Eos+ inter- and intra-ray fibroblasts at 4 dpa (Figure 2E-G). These inter- and intra-ray fibroblasts were part of continuous, converted Eos-expressing cell populations extending to the lateral and medial distal blastema, respectively. These results further support blastemal mesenchyme as a common source for transcriptionally similar intra- and inter-ray fibroblasts, while suggesting spatially segregated re-differentiation paths.

### Insulin-like growth factor / mTOR signaling elevates global translation in mixed-lineage blastema cells

We next aimed to identify candidate processes regulating mesenchyme and osteoblast state transitions during regenerative outgrowth. We first evaluated Gene Ontogeny terms associated with DEGs across mesenchyme/fibroblast pseudotime, identifying overrepresentation of protein translation associated terms in blastemal mesenchyme compared to fibroblasts (Figure 3A). Components of IGFR/mTOR signaling, which widely promotes translation (reviewed in Thoreen, 2017), were upregulated in blastemal mesenchyme vs. fibroblasts transcriptomes (Figure 3B), matching a published report (Hirose et al., 2014). Chemically inhibiting mTOR or IGF receptor signaling at 3 dpa confirmed Insulin/IGF promotes mTOR-dependent ribosomal S6 kinase activity in blastemal mesenchyme and osteoblasts (Hirose et al., 2014; Figure Supplement 3-1). We next used O-propargyl-puromycin (OPP), a stainable puromycin analog incorporated during translation (Liu et al., 2012), to identify dramatically elevated bulk translation in blastemal mesenchyme and osteoblasts at 3 dpa. In contrast, translation levels were much lower in the most distal blastema, epidermis, and proximal regenerated tissue (Figure 3C, D). mTOR inhibition with Torin at 3 dpa decreased OPP incorporation across cell types, confirming IGFR/mTOR specifically elevates translation in blastemal mesenchyme and osteoblasts during regenerative outgrowth (Figure 3E, F).

**Figure 3.**
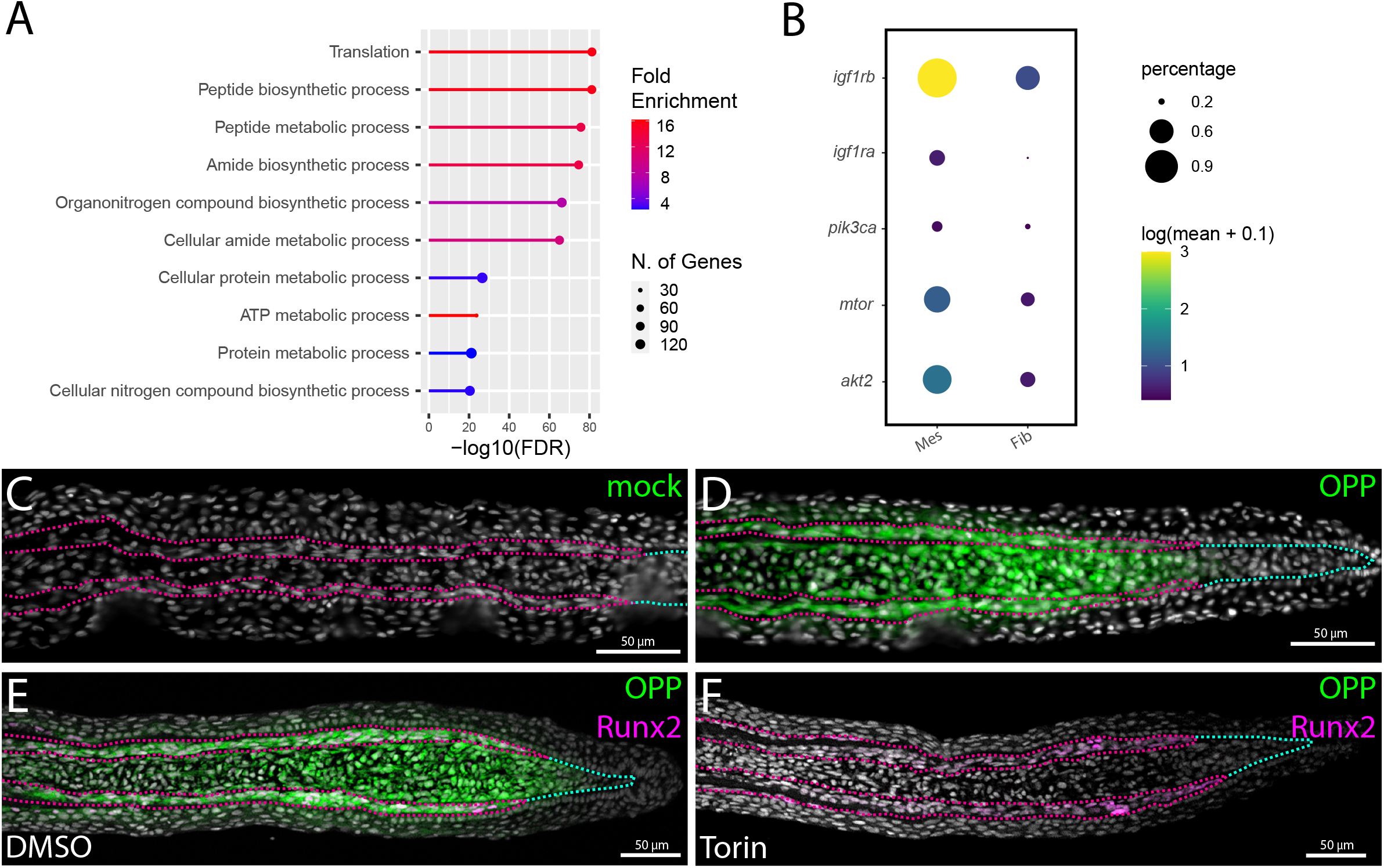
IGFR/mTOR signaling greatly elevates bulk translation in blastemal mesenchyme and differentiating osteoblasts during regenerative outgrowth. (A) Biological process Gene Ontogeny (GO) terms overrepresented in genes differentially expressed across pseudotime in fibroblast lineage cells. Terms associated with protein translation are enriched in blastemal mesenchyme (red) relative to fibroblasts (blue). The size of the circle indicates the number of enriched transcripts defining a given GO term. (B) Gene list and individual expression metrics for differentially expressed insulin growth factor and mTOR pathway components in fibroblast lineage cells. Mes: mesenchyme, fib: fibroblast. The color indicates relative expression levels and relative circle size reflects the percentage of cells expressing the given transcript. (C, D) Confocal images of regenerating fin sections (3 day post amputation (dpa)) stained for O-propargyl-puromycin (OPP)-labeled nascent translation (green) after a 30 minute mock (C) or OPP (D) exposure immediately prior to collection. (E, F) Confocal images of 3 dpa fin sections from DMSO control or Torin mTOR inhibitor-treated fish (during the two hours prior to harvest), stained for OPP (green) and Runx2 osteoblast marker (magenta). mTOR inhibition markedly decreases nascent protein production within osteoblast lineage (Runx2+) and adjacent blastemal mesenchyme. Hoechst-stained nuclei are in grey. Magenta dashed lines outline osteoblasts. White dashed lines indicate distal blastema with relatively low OPP labeling. All scale bars are 50 *μ*M.

### IGFR/mTOR accelerates fin osteoblast maturation *in vitro* and regeneration *in vivo*

IGFR/mTOR may support both osteoblast proliferation and differentiation during the fin regeneration outgrowth phase (Hirose et al., 2014). However, we found IGF-1 was insufficient to differentiate zebrafish fin pre-osteoblast AB.9 cells (result replicated within Figure 4A-C; AB.9 cells characterized in Paw and Zon, 1999, Yette et al., 2021). We profiled differentially expressed ligand receptors across osteoblast UMAP space to identify candidate pathways that cooperatively could facilitate pOb maturation (Figure 4A, select transcripts in Figure 4B). We discounted BMP receptors, previously implicated in fin osteoblast differentiation (Stewart et al., 2014), given AB.9 cells autonomously activated BMP signaling without differentiating (Figure Supplement 4-1). We then focused on pathways previously associated with osteoblast maturation (reviewed in Plotkin and Bruzzaniti, 2019, Mangiavini et al., 2022, Hacemi et al., 2018). We assessed the ability of corresponding ligands (IGF-1, FGF, EGF, PDGF), a canonical Wnt agonist (the GSK3 inhibitor CHIR99021), and a glucocorticoid receptor agonist (Dexamethasone (Dex)) to promote AB.9 cell differentiation. We examined elevated alkaline phosphatase activity (Quarles et al., 1992), transition from mesenchymal to epithelial state (Stewart et al., 2014), and decreased cell density indicative of reduced proliferation after four days treatment with the combined activators or each individual factor together with IGF-1. Only dexamethasone (Dex), an agonist of glucocorticoid receptor signaling, induced AB.9 cell maturation by all criteria. Unexpectedly, Dex robustly promoted differentiation even in the absence of IGF-1 (Figure 4C-E).

**Figure 4.**
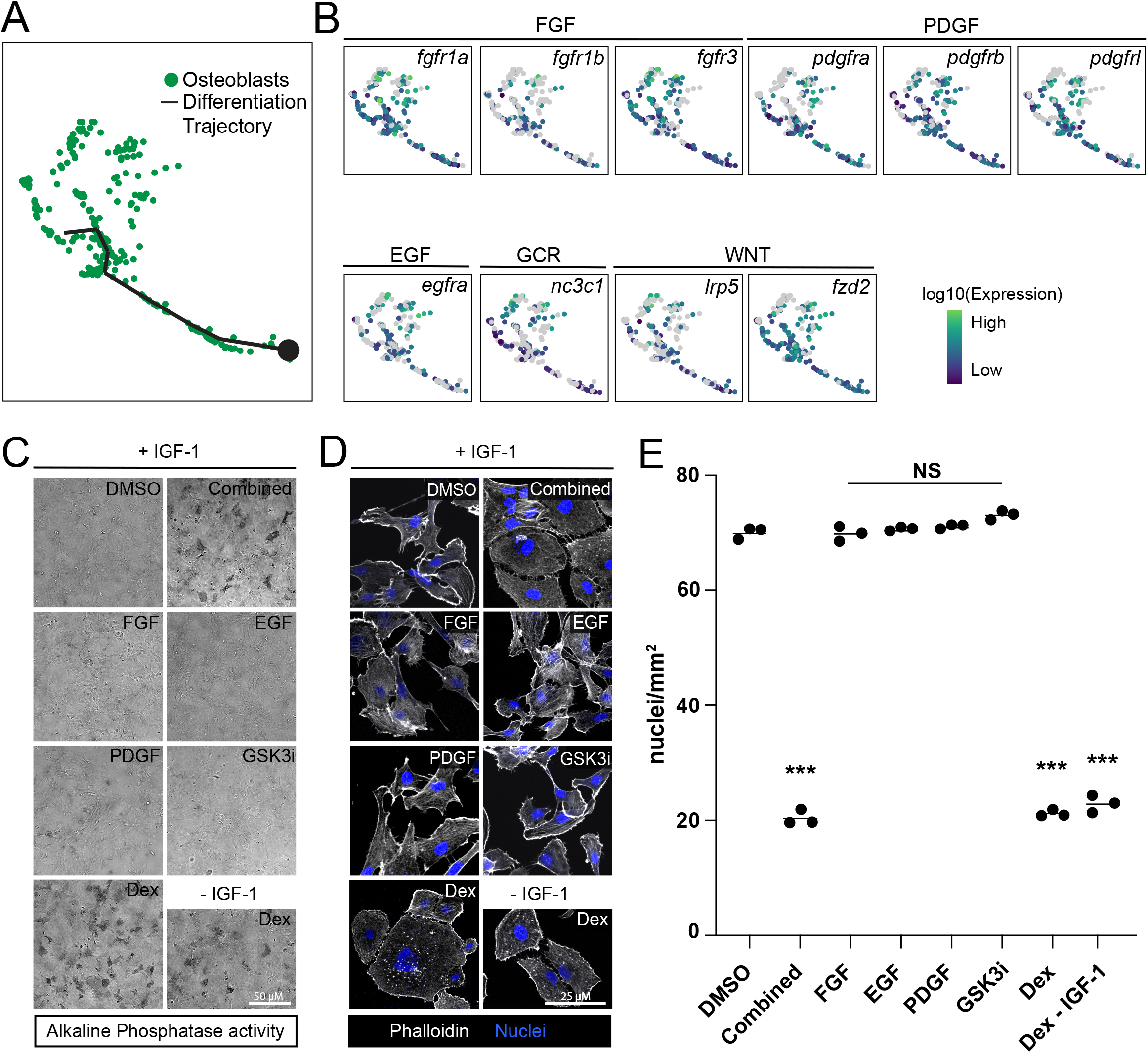
Exogenous glucocorticoids promote fin osteoblast differentiation in vitro. (A) UMAP plot of the defined osteoblasts from the outgrowth phase scRNA-Seq dataset with an overlaid pseudotime trajectory. (B) Osteoblast UMAP plots for indicated transcripts encoding receptors associated with FGF, PDGF, EGF, Glucocorticoid (GCR) and Wnt signaling. (C) Brightfield images of AB.9 fin pre-osteoblasts treated for 4 days with IGF-1 (except bottom right panel) and candidate osteoblast differentiation factors (recombinant ligands or small molecules) and then stained for alkaline phosphatase activity. (D) Fluorescent confocal images of same AB.9 groups from (C) stained for phalloidin (white) and nuclei (Hoechst, magenta). (E) Data point graph showing nuclei per mm^2^ of AB.9 cell culture groups from (D). Triplicate samples with means are shown. Dexamethasone (Dex), a glucocorticoid receptor agonist, is sufficient to induce osteoblast maturation as indicated by alkaline phosphatase activity, epithelial-like actin cystoskeleton organization, and reduced proliferation implied by decreased nuclei density. ***: p < 0.001, NS: p > 0.05.

Dex sufficiency to induce osteoblast maturation *in vitro* suggested IGFR/mTOR-driven bulk protein translation may accelerate rather than instruct osteoblast differentiation. We tested this hypothesis by incubating AB.9 pObs with OPP while stimulating IGF and/or glucocorticoid signaling. IGF-1 caused elevated translation independent of Dex addition (Figure 5A, B). Neither IGF-1 nor Dex altered global nascent transcription as monitored by 5-ethynyl uridine (5-EU) incorporation (Figure 5A, B). Therefore, IGF-1 increases global translation in AB.9 cells irrespective of glucocorticoid activity. We then directly tested if IGF-1 accelerates Ob maturation. Combined IGF and Dex induced widespread AB.9 cell epithelialization within one day, whereas Dex alone required at least three days (Figure 5C). Consistently, in vivo mTOR inhibition during the outgrowth phase by Torin exposure from 3 to 14 dpa slowed but did not prevent continued fin regeneration, including formation of segmented bony rays (Figure 5D, E).

**Figure 5.**
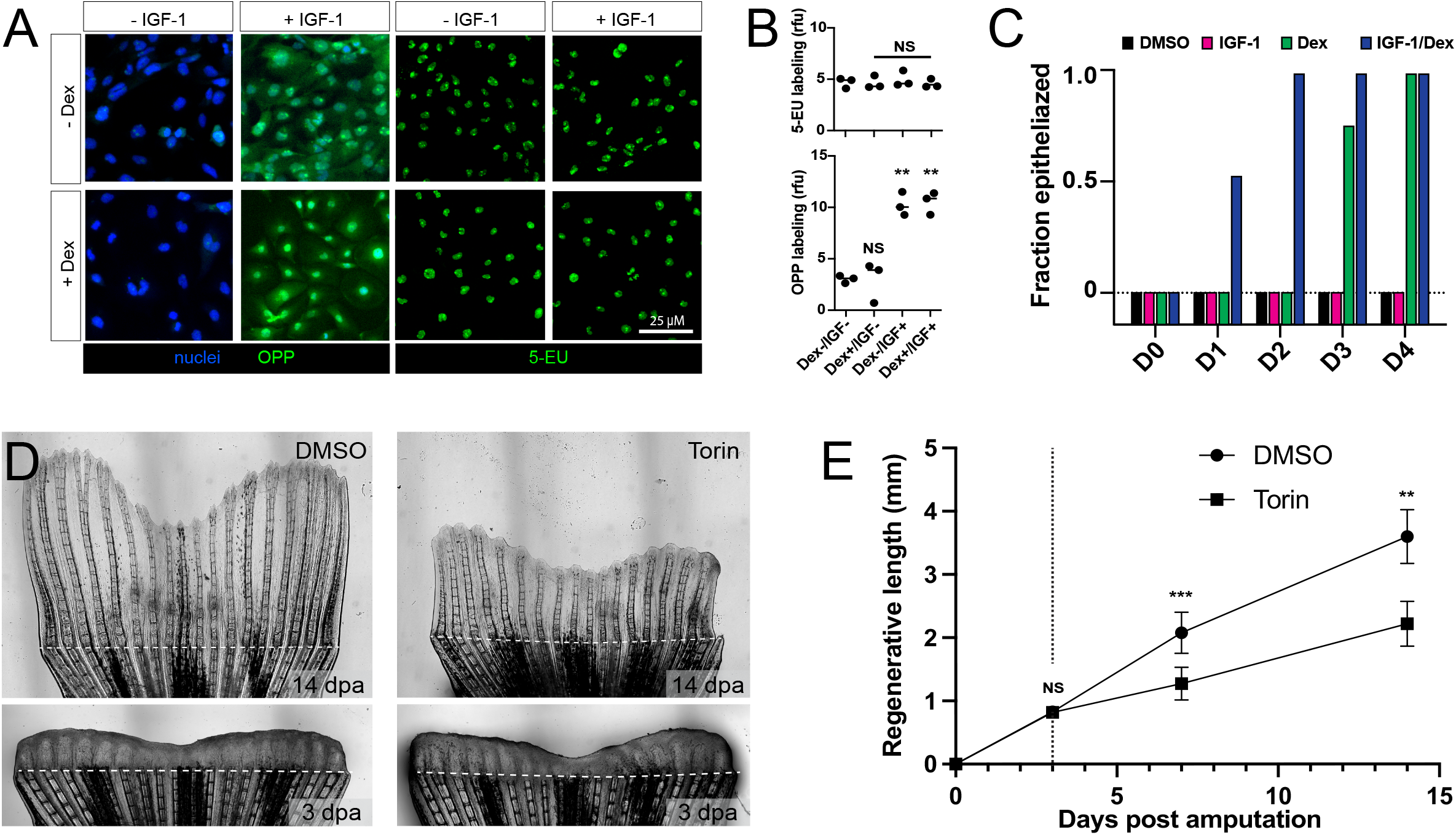
IGFR/mTOR signaling amplifies translation to accelerate osteoblast differentiation and regenerative tempo. (A) Confocal fluorescent images of AB.9 pre-osteoblasts treated with or without IGF-1 and/or dexamethasone (Dex) for 48 hours and stained for O-propargyl-puromycin (OPP, left panels, green) or 5-ethynyl uridine (5-EU, right panels, green) incorporation. IGF-1 stimulates elevated translation independent of Dex. Neither IGF and/nor Dex stimulation alter global transcription as indicated by 5-EU incorporation. Scale bar is 25 *μ*M. (B) Individual data point plots scoring intensity of 5-EU or OPP incorporation in cell groups shown in (A). Means are shown. **: p < 0.01, NS: p > 0.05. (C) Bar graphs showing the fraction of AB.9 plates (n=6, from two independent experiments) with a phalloidin-monitored epithelial sheet-like organization at zero through four days (D0-D4) after DMSO, IGF-1, Dex, or IGF-1/Dex treatments. IGF-1/Dex1 accelerated AB.9 differentiation relative to Dex alone as indicated by the earlier induction of epithelial characteristics. (D) Differential interference contrast stitched images of regenerating caudal fins treated with DMSO or the mTOR inhibitor Torin starting at 3 days post amputation (dpa). Images are acquired at 3 and 14 dpa. The dashed line marks the amputation position. (E) Line graphs showing mean regenerated caudal fin lengths of DMSO and Torin-treated fish from animals shown in (D). Error bars are one standard deviation. The dashed vertical line shows the onset of drug treatment at 3 dpa. Fin regeneration proceeds at a slower rate but still produces segmented bony rays in spite of continuous mTOR inhibition. **: p < 0.01, ***: p < 0.001, NS: p > 0.05.

## DISCUSSION

We characterize a new single cell transcriptome dataset focused on fibroblast/osteoblast lineages during the zebrafish fin regeneration outgrowth phase. We validate *robo3* as a novel marker of the “organizing center” or “niche” growth-promoting and fibroblast-lineage distal blastema cells. In contrast, *fhl1a* defines proximal intra-ray fibroblasts. We further identify *rgs5a* versus *slit2* as distinguishing markers of inter- and intra-ray fibroblasts, respectively, which otherwise share highly similar transcriptomes. Several studies show regenerated intra-ray fibroblasts differentiate from distal blastemal mesenchyme (Tu et al., 2011, Stewart et al., 2012, Tornini et al., 2016). A pseudotemporal trajectory and photoconvertible lineage tracing indicate inter-ray fibroblasts also derive from blastema mesenchyme, likely from cells at the lateral aspects of the far distal blastema. These cells seemingly migrate bi-laterally distal to ray-forming pre-osteoblasts and then organize into a fibroblast network that supports inter-ray tissue. Meanwhile, intra-ray fibroblasts originate from relatively central and proximal blastemal mesenchyme that is encompassed by pre-osteoblasts. However, our studies do not resolve if intra- and inter-ray fibroblasts are pre-specified within distal blastema mesenchyme. More extensive single cell profiling and cell-refined lineage tracing would help resolve spatially segregated fibroblast-lineage populations and their state transitions within a compartmentalized blastema.

We profiled pseudotime trajectories to identify that IGFR/mTOR elevates global protein translation in re-differentiating mesenchyme and osteoblasts, consistent with a previous report (Hirose et al., 2014). We used cultured fin pre-osteoblasts to show that mTOR accelerates osteoblast maturation. Consistently, IGFR/mTOR inhibition during the outgrowth phase reduced the rate of fin regeneration without affecting bone deposition. We propose that, during outgrowth, IGFR-driven mTOR activity elevates translation to coordinate cellular outputs across lineages and thereby modulate regenerative tempo.

In mammals, endogenous glucocorticoids promote bone formation whereas exogenous glucocorticoids cause osteoporosis, including by direct effects on osteoblasts that may be pro- or anti-differentiation dependent on osteoblast progenitor state (Reviewed in Lee and Tuckerman, 2022). Prolonged glucocorticoid stimulation also causes osteoporosis in adult zebrafish fin rays (Bohns et al., 2020). Further, elevating glucocorticoids during fin regeneration reduces ray length and disrupts osteoblast function (Geurtzen et al., 2017, Schmidt et al., 2019). These gain-of-function phenotypes could reflect secondary effects, for example by precocious osteoblast differentiation depleting proliferative progenitor osteoblasts. Regardless, potential in vivo roles of endogenous glucocorticoid signaling during fin regeneration remain untested. The AB.9 fin pre-osteoblast cell line, initially described as fibroblast-like (Paw and Zon, 1999), provides an avenue for future mechanistic studies of glucocorticoid and IGFR/mTOR signaling in osteoblasts, including comparisons with IGFR/mTOR pro-differentiation roles in mammalian osteoblasts (Chen et al., 2014, Xian et al., 2012). Evaluating IGFR/mTOR effects on fin fibroblast lineage state transitions would benefit from similar cell culture tools.

IGFR/mTOR signaling also is required for the initial fin injury response (Hirose et al., 2014). This is a seemingly essential, switch-like rather than modulatory role. Nonetheless, IGF/mTOR’s establishment and outgrowth roles could share a common mechanism in broadly elevated protein translation. Intriguingly, axolotl limb resection also triggers a rapid increase in mTOR-dependent protein activation essential to initiate regeneration (Zhulyn et al., 2021). This effect is most pronounced in epidermal cells, consistent with the early role of the wound epidermis in triggering a regenerative response. Likewise, IGF/mTOR-dependent S6K activity is high in epidermal cells of establishing but not outgrowing fin regenerates (Hirose et al., 2014). Therefore, mTOR may be dynamically activated in distinct cell lineages throughout regeneration depending on the biological processes (e.g. de-differentiation, migration, proliferation, re-differentiation) active at each regeneration stage.

Igf2b may have local, essential roles in fin blastema establishment and regenerative outgrowth (Chablais and Jazwinska, 2010). Insulinopenia and hyperglycemia upon streptozocin damage to pancreatic beta cells reduces fin regeneration (Olsen et al., 2010, Carvalho et al., 2017). Therefore, local IGFs and/or systemic insulin, the latter dependent on a re-established vasculature, could be responsible for the observed IGFR- and mTOR-dependent protein translation and downstream tempo modulation. A potential systemic contribution suggests how the overall regeneration rate could be continuously modulated to suit the animal’s physiologic state and environment. For example, fin regeneration rate, but not ultimate regenerated fin size, correlates with water temperature (Johnson and Weston, 1995). Nutrient availability manifesting in elevated circulating insulin and/or direct cell metabolic effects on mTOR activity could also promote faster regeneration. In support, leucine and glutamine treatment elevate mTOR activity and cell proliferation at least during the fin blastema establishment phase (Takayama et al., 2018). In *Xenopus tropicalis*, nutrient deprivation likely acts through inhibited mTOR to curtail tail regeneration during the larval refractory period (Williams et al., 2021). Therefore, regulated mTOR may serve as an evolutionarily-conserved rheostat to modulate the tempo of regeneration to suit environmental conditions without altering the restored organ form. By extension, enhancing mTOR-dependent translation may be a promising approach to augment human injury repair responses without compromising orthogonal pattern-restoring networks.

## MATERIALS AND METHODS

### Zebrafish husbandry and fin amputations

Zebrafish were maintained at ∼28.5°C under 14:10 light:dark cycles. The following established lines were used: wildtype (ABC), *Tg(sp7:EGFP)*^*b1212*^ (DeLaurier et al., 2010), *Tg(RUNX2:mCherry)* (Barske *et al*., 2020), *Tg(tph1b:mCherry)* (Tornini *et. al*., 2015), *Tg(sost:nlsEos)* (Thomas and Raible, 2019). Transgenic lines were maintained in the ABC strain background. Adult fish were anesthetized with MS222 and their caudal fins amputated using a razor blade. The animals were then returned to circulating fish water. Euthanasia was performed via MS222 overdose. The University of Oregon Institutional Animal Care and Use Committee (IACUC) approved and monitored all zebrafish procedures following the guidelines and recommendations in the Guide for the Care and Use of Laboratory Animals (National Academic Press).

### Confocal imaging

Images were acquired with a Nikon Eclipse T*i-*E widefield microscope equipped with a Yokogawa CSU-W1 spinning disk confocal scanner unit or with a Zeiss LSM 880 laser scanning confocal microscope. Confocal image stacks were processed using ImageJ software to generate single optical slice digital sections and/or maximum intensity projections across multiple channels. Adobe Photoshop was used to adjust display levels with identical processing settings for a given experiment. Figure panels were compiled in Adobe Illustrator.

### Eos photoconversion and imaging

*Tg(sost:nlsEos)* fish were anesthetized as described above. Fins were viewed on a Nikon Eclipse T*i-*E microscope with a Yokogawa CSU-W1 spinning disk confocal scanner unit. Eos-expressing cells of the blastema were photoconverted with 405 nm laser illumination for 1 minute. Before and after images were acquired to confirm photoconversion of Eos from green (518 nm) to red (580 nm) emission. Fish were returned to system water and then similarly re-imaged after defined periods.

### Paraffin section immunostaining

Amputated caudal fins were fixed in 4% paraformaldehyde (PFA)/PBS overnight at 4°C, processed for paraffin sections, sectioned, and stained largely as described (Stewart et al., 2014). After PBS washing, fins were decalcified for 4 days in 0.5M EDTA, pH 8.0 with daily solution changes. Fins were dehydrated through an ethanol series, cleared with xylenes, and embedded in paraffin wax. 7 µm sections were obtained with a Leica RM255 microtome. Antigen retrieval was performed on rehydrated sections using 1 mM EDTA + 0.1% Tween-20 or 0.05% citraconic anhydride for 5 minutes in a pressure cooker. After PBS washes, sections were blocked in 1x PBS with 10% nonfat dry milk alone or 10% nonfat dry milk with 2% normal goat serum and 4% fetal bovine serum for a minimum of 1 hour. Primary antibodies were incubated overnight at 4°C in blocking solution (see key reagents table). Sections were washed in PBS containing 500 mM NaCl + 0.1% Tween-20. Alexa Fluor conjugated secondary antibodies (Thermo Fisher) were diluted 1:1000 in blocking buffer and applied to sections for 1 hour at room temperature. Sections were washed, nuclei labeled with 1:1500 Hoechst dilution (stock 10 mg/mL), and mounted in SlowFade Gold Antifade (Thermo Fisher).

### Cryosectioning and staining

Fin tissue was collected into fresh 4% PFA in 1x PBS on ice. Tissue was fixed at 4°C for 4 hours, then washed with cold 1x PBS three times, 5 mins each. PBS was replaced with sterile 30% sucrose in 1xPBS (cold) and left on a rocker overnight at 4°C. Fin tissue was then embedded in Tissue-Tek O.C.T. compound (Sakura Finetek) and frozen on dry ice. Tissue was stored at -80°C until ready to section. 12 *μ*m thickness cryosections were collected on slides, dried and stored at -20°C. Immunofluorescence staining was performed using the zns-5 antibody (ZIRC) at a concentration of 1:200 in a 10% milk, 1x PBS, 0.1% Tween-20 blocking solution. Alexa Fluor conjugated secondary antibody was used at 1:500 and nuclei labeled with 1:1500 Hoechst dilution (stock 10 mg/mL). Slides were mounted using SlowFade Diamond Antifade (Thermo Fisher).

### RNAscope in situ hybridization

RNAscope ((Wang et al., 2012) probes for *rgs5a, slit2*, and *robo3* were designed and synthesized by ACD Bio and visualization performed using the Multiplex Fluorescent kit (ACD Bio) according to the manufacturer’s recommendations for paraffin wax embedded sections. Combined immunostaining was carried out after RNAscope, as directed by the manufacturer (ACD Bio). Nuclei were visualized by Hoechst staining and slides mounted as above (Thermo Fisher).

### *In vivo* drug treatment and O-propargyl-puromycin labeling

O-propargyl-puromycin (OPP, Thermo Fisher) was dissolved in DMSO at 2 mg/ml stock and frozen at -20°C. Immediately prior to injection, OPP was thawed and diluted 1:10 in injection buffer (50% PEG-400, 5% propylene glycol, 0.5% Tween 80). 10 *μ*l of this solution was injected intraperitoneally (2 mg/kg) into regenerating animals 2 hours prior to tissue collection. Where indicated, cohorts were pre-treated for 2 hours in mTOR inhibitor (Torin, final concentration 100 nM in fish water, Cayman Chemical). For experiments using rapamycin (LC Labs) to inhibit mTOR, the drug was dissolved in DMSO, diluted 1:10 in injection buffer (see above) and injected intraperitoneally at a dose of 10 mg/kg. To block IGFR signaling, animals were transferred to tanks containing 5 *μ*M IGFR inhibitor (NVD-ADW742, MedChemExpress). Rapamycin and NVD- ADW742 treatments were from 64-72 hpa.

### Cell culture

AB.9 cells (Paw & Zon, 1999) were purchased from ATCC (CRL-2298) and cultured in Dulbecco’s Modified Eagle’s Medium (DMEM) supplemented with fetal bovine serum (10%) and 1X penicillin-streptomycin (Thermo Fisher) at 30°C in 5% CO_2_ and 5% O_2_. AB.9 subclones were first isolated by plating cells at clonal dilution in a 96 well plate. Colonies containing cells with uniform morphology and robust growth were picked and stained with Runx2 and Sp7 antibodies (Stewart et al., 2014, Yette et al., 2021) to confirm pre-osteoblast state. Cells of these subclones were Runx2^+^ with low or absent Sp7 protein. One of these clones, AB.9.8, was expanded and used for the experiments shown. For differentiation assays, candidate growth factors were reconstituted in 1x PBS or water and used at a final concentration of 100 ng/mL. Dexamethasone (final concentration 100 nM) and GSK3 inhibitor CHIR99021 (final concentration 5 μM) were dissolved in DMSO. Media was removed, cells washed, and media replenished every 24 hours during differentiation assays.

### Alkaline phosphatase assay

AB9.8 cells were washed once with PBS, and then fixed for 1 minute with 4% PFA in PBS. Fixed cells or fins were washed once with PBS and once more with PBS containing 0.1% Tween-20. Cells or fins were washed once for 5 minutes in developing buffer (100 mM Tris pH 9.5, 100 mM NaCl, 10 mM MgCl_2_) and then developed in the dark with fresh developing buffer supplemented with 225 *μ*g/ml nitro blue tetrazolium (NBT) (Promega) and 175 *μ*g/ml 5-bromo-4-chloro-3- indolyl-phosphate (BCIP) (Promega). Reactions were stopped as the signal became apparent with three PBS rinses. Cells or fins were fixed with 4% PFA in PBS for 5 minutes at room temperature and then washed three times for 5 minutes.

### Phalloidin assay

AB.9 cells were washed once with PBS then incubated in PBS containing 0.1% Tween-20 + 2% PFA for 30 minutes. Cells were washed 3 times in PBS then incubated in 1:40 phalloidin (Thermo Fisher) in PBS for 20 minutes at room temperature in the dark. Cells were washed 3 times with PBS to remove excess stain. Nuclei were labeled using a 1:1500 Hoechst dilution (stock 10 mg/mL) prior to imaging. For the time course experiment, individual plates (six for each condition from two experimental replicates) were scored as having predominantly epithelial (interconnected cells with plasma membrane-associated actin filaments) or mesenchymal characteristics.

### Fin length and cell count measurements

Caudal fin lengths and cell counts were measured using ImageJ-Fiji (National Institutes of Health). Fin lengths were defined as the length of the third dorsal ray starting from the amputation plane and measured using the measure tool. Cells were identified by the presence of Hoechst-labeled nuclei and counts were determined using the cell counter tool. Fin length and cell count statistical comparisons used Student’s two-tailed *t*-tests with two-stage step-up method of Benjamini, Krieger and Yekutieli for multiple comparisons and one-way ANOVA with Tukey post-hoc tests, respectively.

### *In vitro* O-propargyl-puromycin (OPP) and 5-ethynyl uridine (5-EU) labeling

Cells were incubated in either 20 mM O-propargyl-puromycin (OPP) (Thermo Fisher) or 0.2 mM 5-ethynyl uridine (5-EU) (Thermo Fisher) in media for 30 minutes according to kit instructions. Samples were then fixed using 4% PFA in PBS followed by permeabilization using 0.5% Triton X-100 for 10 minutes. OPP visualization used Click-iT OPP detection followed by HCS NuclearMask Blue Stain (Thermo Fisher). 5-EU visualization used hapten-containing azide and Click-iT Cell Reaction Buffer Kit (Thermo Fisher). Samples were washed twice before imaging. Plates were imaged with a Nikon Eclipse T*i-*E microscope with a Yokogawa CSU-W1 spinning disk confocal scanner unit. The ImageJ-Fiji measure tool was used to determine relative florescence intensities (RFU). RFU statistical comparisons used one-way ANOVA with Tukey’s post-hoc tests.

### Single cell collection, scRNA-Seq library construction and sequencing

Regenerated tissue distal to the amputation plane from *Tg(tph1b:mCherry);Tg(sp7:egfp)* animals at 3 and 7 days post amputation was collected, pooled, and dissociated. Tissue from 20 animals was pooled for each group. Regenerative fin tissue was collected into 1x PBS on ice, diced, then washed with L-15 media supplemented with 1X GlutaMax, 1X Antibiotic-Antimycotic and 1X Penicillin-Streptomycin (all Thermo Fisher). The tissue was then enzymatically digested using 0.25% Trypsin-EDTA (Thermo Fisher) and Liberase DL (2 mg/ml, Sigma) at room temperature for 20 min followed by mechanical dissociation by pipet titration every 5 minutes. Enzymatic digestion was inhibited by the addition of 20% fetal bovine serum. Cells were filtered through a 100 μm cell strainer and then pelleted by centrifugation at 0.2 g for 2.5 minutes. The cell pellet was washed with PBS and re-suspended in PBS + 0.04% bovine serum albumin. For each sample, ∼5000 cells were inputted to the Chromium platform (10X Genomics) with samples split evenly between two lanes each. Single-cell mRNA libraries were prepared using the single-cell 3’ solution V3 kit (10X Genomics). Quality control and quantification assays were performed using a Qubit fluorometer (Thermo Fisher) and a Fragment Analyzer (Agilent). Libraries were sequenced with an NextSeq 500 (llumina) using 75-cycle, high output kits (read 1: 26 cycles, i7 Index: eight cycles, read 2: 57 cycles). Each sample was sequenced to an average depth of 240 million total reads. Based on capture rate, this resulted in an average read depth of ∼30,000 reads/cell after read-depth normalization.

### scRNA-Seq data processing

Cell barcodes, UMIs, and cDNA reads were input into Kallisto version 0.46.1 (Bray et al., 2016) configured for 10x v3 chemistry. Pseudoalignment used a transcriptome generated from zebrafish GRCz11. Bustools version 0.39.3 (Melstead et al., 2019) was used for sequence correction, counting and matrix processing to remove cells expressing less than 150 genes and disregard genes expressed in less than 20 cells. Monocle3 version 0.2.3 (Cao et al., 2019) was used to generate a cell-data-set file.

### UMAP visualization and clustering

Uniform Manifold Approximation and Projection (UMAP) (McInnes et al., 2018) was used to project cells in two dimensions. Community detection clustering (Levine, et al., 2015) used the reduceDimension and clusterCells functions in Monocle3 (v.3.0.2.3) with default parameters (except for, reduceDimension: reduction_method = UMAP). Clusters were assigned as particular cell types by manual perusal of enriched genes for established marker genes. Cells isolated from 3 dpa and 7 dpa animals were combined to maintain consistent analysis across stages of regenerative outgrowth. Batch correction was used between the two groups to avoid sample-specific separation or clustering in UMAP space. Cells representing the top 5% of UMIs were discarded as potential doublets (predicted frequency of ∼2-3% based on 5000 cell input). All genes were used for Principal Components Analysis (PCA). The top 20 principal components (based on the generated scree plot) were used for UMAP projection. To subcluster the fibroblast/osteoblast cluster, identified osteoblasts were removed and a community detection clustering re-applied to the remaining putative fibroblast/mesenchyme cells.

### Differential gene expression, pseudotime and Gene Ontogeny (GO) analysis

Differentially expressed genes in subclusters were identified using the principalGraphTest function in Monocle3 (v.0.2.3) (neighbor_graph = knn) with default parameters (Cao et al., 2019). UMAP dimensionality reduction and community detection clustering was re-applied to fibroblast/mesenchyme cells and a pseudotemporal trajectory was defined using learn_graph and order_cells in Monocle3 using default parameters. The pseudotime root was set within actively cycling cells of the blastemal mesenchyme subcluster 1 to continue through fibroblast UMAP space. The principalGraphTest function in Monocle3 (v.0.2.3) across pseudotime (neighbor_graph = principal_graph) with default parameters (Cao et al., 2019) identified genes expressed for gene ontology term analysis. Genes with Moran’s I greater than 0.5 were subset for Gene Ontology (GO) analysis as those most significantly sequestered to unique pseudotemporal space (Supplemental File 3). shinyGO v0.75 was used for a statistical overrepresentation test identifying enriched GO biological processes and associated gene sets.

## ACKNOWLEDGEMENTS

We thank the University of Oregon AqACS Facility for zebrafish care; the University of Oregon zebrafish community for support; David Raible for providing *Tg(sost:nlsEos)* fish; and the Stankunas lab for input.

## FOOTNOTES

### Author contributions

V.M.L, H.K.L., S.S. and K.S. designed experiments; V.M.L, H.K.L., A.L.H, H.M., A.E.R., and S.S. performed experiments; V.L. and K.S. prepared and wrote the manuscript with input from all authors.

### Data and material availability

The scRNA-Seq dataset is being deposited to the NCBI Gene Expression Omnibus. Requests for materials should be addressed to K. S.

---

**Figure 1 Supplement 1.**
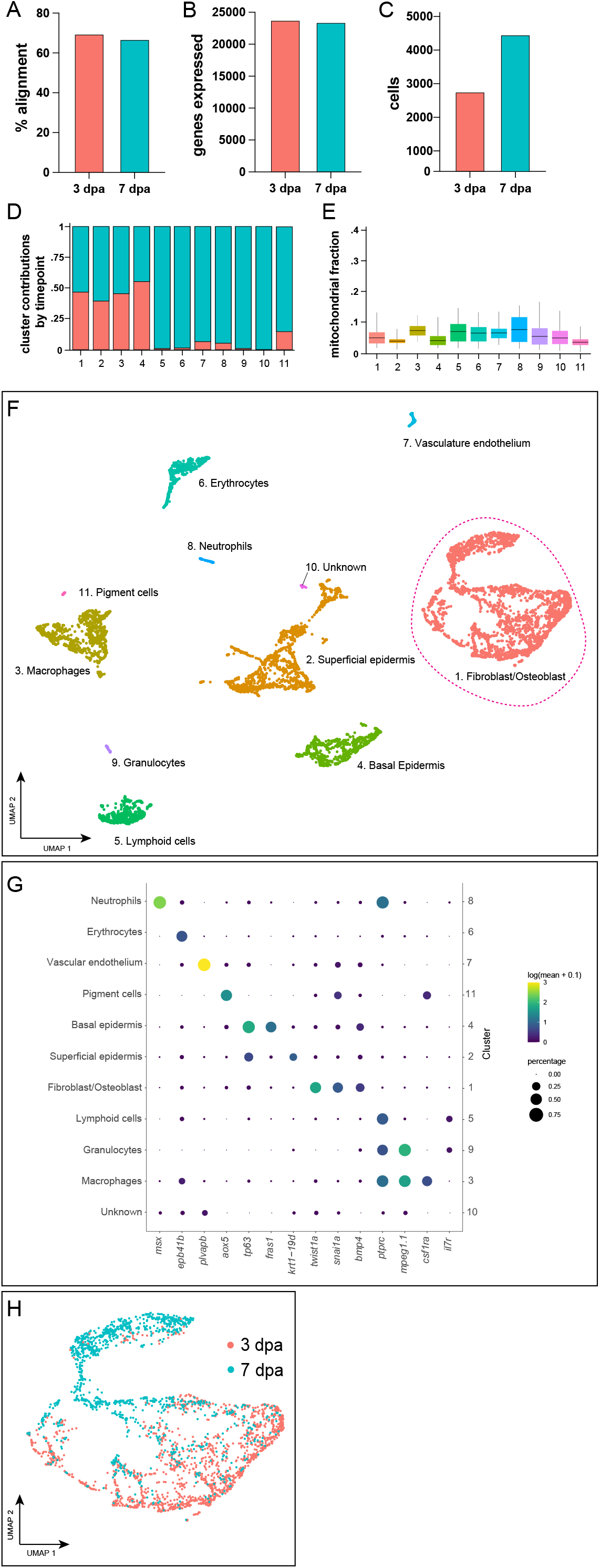
Characteristics of the full scRNA-Seq dataset of fin regeneration outgrowth phases. (A-C) Fraction alignment, number of genes expressed, and number of cells captured by sample used in constructing scRNA-Seq dataset. (D) Fraction cell contribution to individual clusters by sample. (E) Mitochondrial read fraction by cell type following quality control. Colors correspond to cell types labeled in UMAP plot, and graph shows medians with boxes spanning interquartile range and vertical lines indicating farthest extent of data. (F) 2-D UMAP space showing clustering of cell types identified in the scRNA-Seq dataset. Fibroblast/osteoblast lineage cluster outlined with dashed magenta line. (G) Established lineage-specific markers identify cell types within the scRNA-Seq dataset: co-clustered fibroblast and osteoblast cells [cluster 1; *twist1a+, snai1a+, bmp4+*], superficial epidermis [cluster 2; *krt4+*], macrophages [cluster 3; *mpeg1*.*1+*], basal epidermis [cluster 4; *tp63+, fras1+*], lymphoid cells [cluster 5; *ptprc+, il7r+, mpeg1*.*1*-], erythrocytes [cluster 6; *epb41b+*], vascular endothelium [cluster 7; *plvapb+*], neutrophils [cluster 8; *ptprc+, msx+*], dendritic cells [cluster 9; *ptprc+, il7r+, mpeg1*.*1*+], a small unidentified cluster [cluster 10], and pigment cells [cluster 11; *aox5+*] (H) UMAP representation of sample specific contribution to fibroblast/osteoblast lineage cluster.

**Figure 1 Supplement 2.**
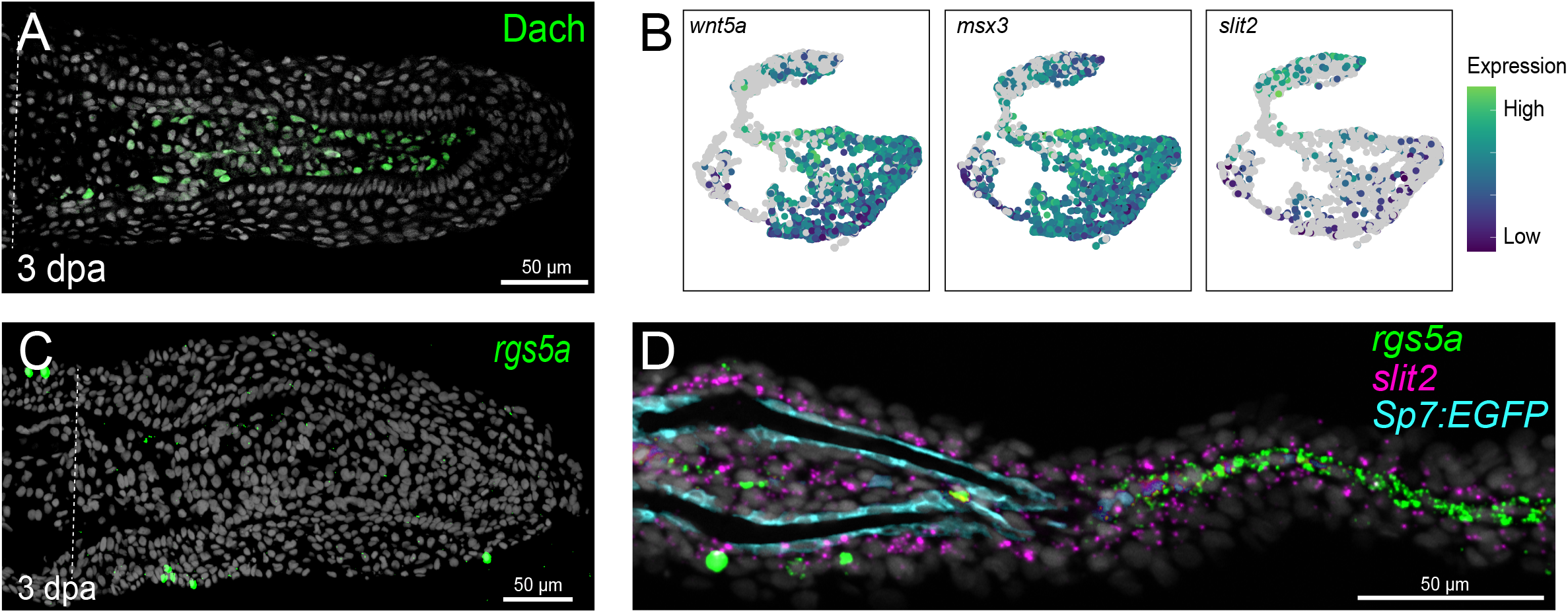
Additional *in vivo* validation of genes marking different populations within fibroblast/osteoblast cluster UMAP space. (A) Dach antibody staining of a 3 dpa fin regenerate section marking distal blastema (or “niche”) cells in green. (B) UMAP plots showing scRNA-Seq dataset expression of known blastema markers *wnt5a and msx3*, and new intra-ray fibroblast marker *slit2* in fibroblast/osteoblast cluster cells during regenerative outgrowth. (C) RNAscope *in situ* hybridization for *rgs5a* (green) in a 3 dpa fin regenerate section shows expression in rare, unidentified superficial epidermal cells but absence within blastemal mesenchyme. (D) RNAscope *in situ* hybridization for *rgs5a* and *slit2* with anti-GFP staining in transverse sections through proximal fin regenerate tissue of a *Tg(sp7:EGFP)* zebrafish. *rgs5a* (green) predominantly marks inter-ray fibroblasts while *slit2* (magenta) expresses in intra-ray fibroblasts and basal epidermal cells. Anti-GFP labeled osteoblasts lining a regenerated bony ray are in cyan. Hoechst-stained nuclei are grey in all section panels. Scale bars are 50 *μ*M.

**Figure 2 Supplement 1.**
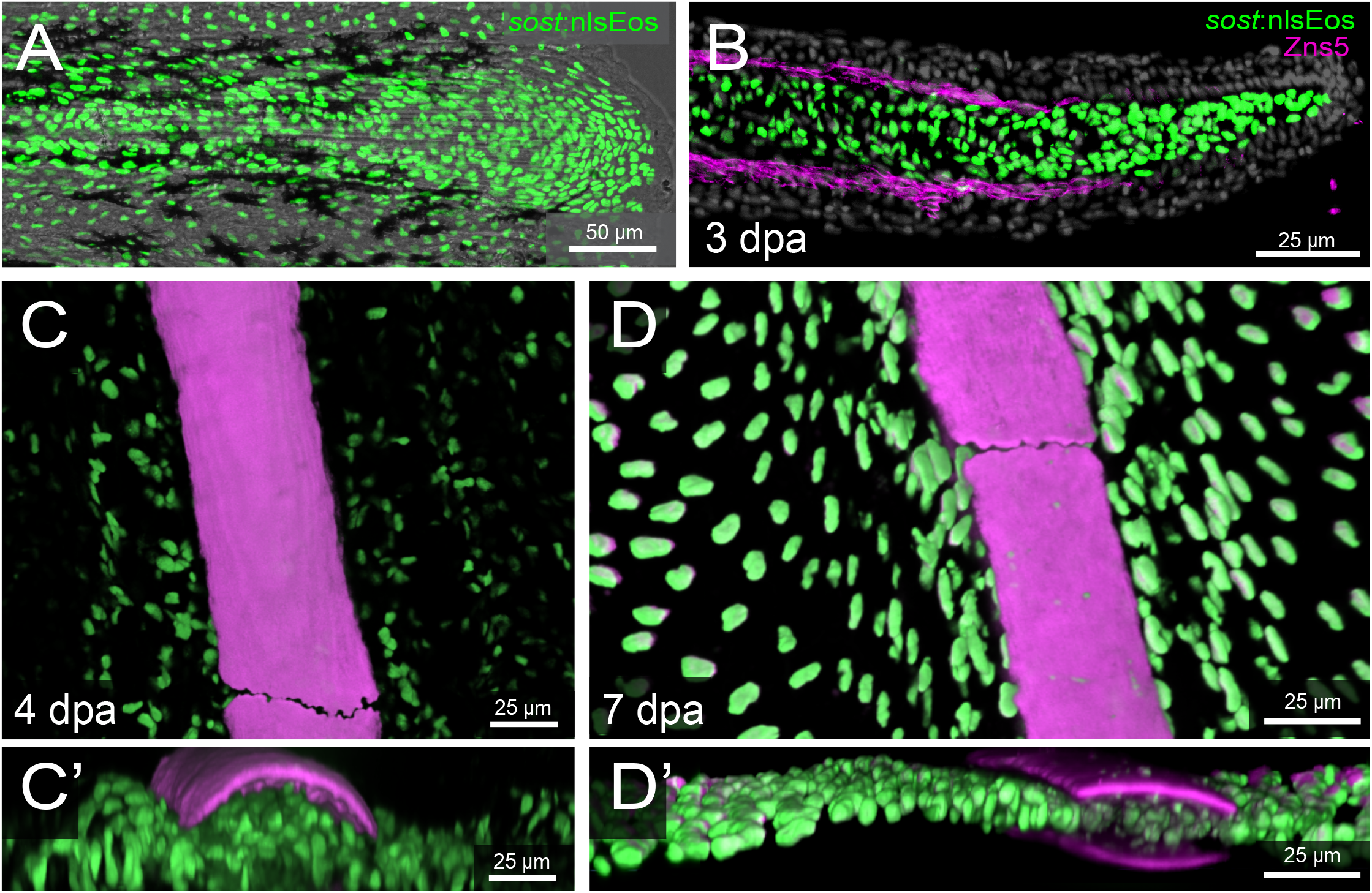
*Tg(sost:nlsEos)* specifically expresses in fibroblast lineage cells of regenerating fins. (A) Maximum intensity projection image of the distal caudal fin of a *Tg(sost:nlsEos)* (referred subsequently as *sost:nlsEos*) fish showing Eos (green) nuclear expression. (B) Antibody stained cryosection of a 3 day post amputation (dpa) 3 dpa *sost:nlsEos* fish showing distinct expression of the osteoblast marker Zns5 (magenta) and Eos (green) within blastemal fibroblast lineage cells. (C-D) 40x confocal reconstructions of whole-mount imaged 4 and 7 dpa fin regenerates shows Eos expression in inter-ray and intra-ray fibroblasts at both outgrowth stages. Eos+ nuclei are green and Alizarin red-stained regenerating bony rays are in magenta. (C’, D’) Transverse view of (C, D). Scales bar are 25 or 50 *μ*M, as indicated.

**Figure 3 Supplement 1.**
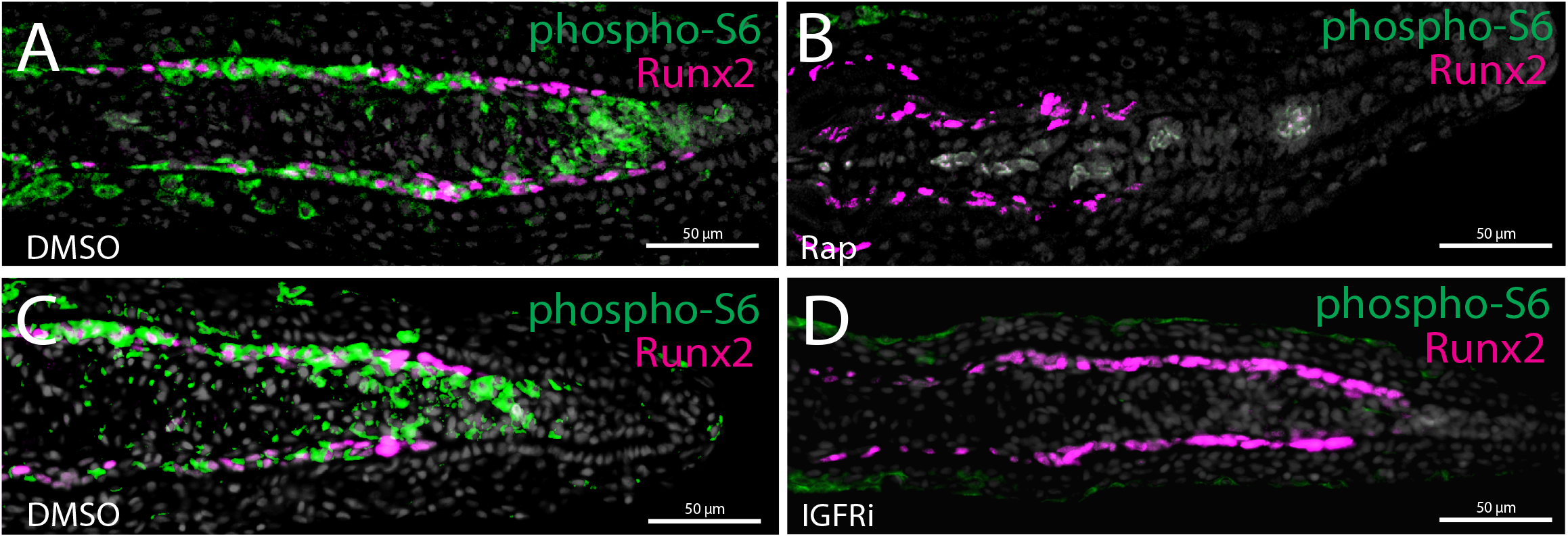
IGF receptor/mTOR-dependent phospho-S6 activity during fin regeneration. (A, B) Decreased phospho-S6 staining (green) of blastemal mesenchyme and osteoblasts (magenta) in mTOR-inhibited (Rap, 6 hours) animals relative to controls (DMSO) at 3 dpa. (C, D) Decreased phospho-S6 staining (green) of blastemal mesenchyme and osteoblasts (magenta) in IGF receptor-inhibited (IGFRi, 6 hours) animals relative to controls (DMSO) at 3 dpa. Scale bars are 50 *μ*M.

**Figure 4 Supplement 1.**
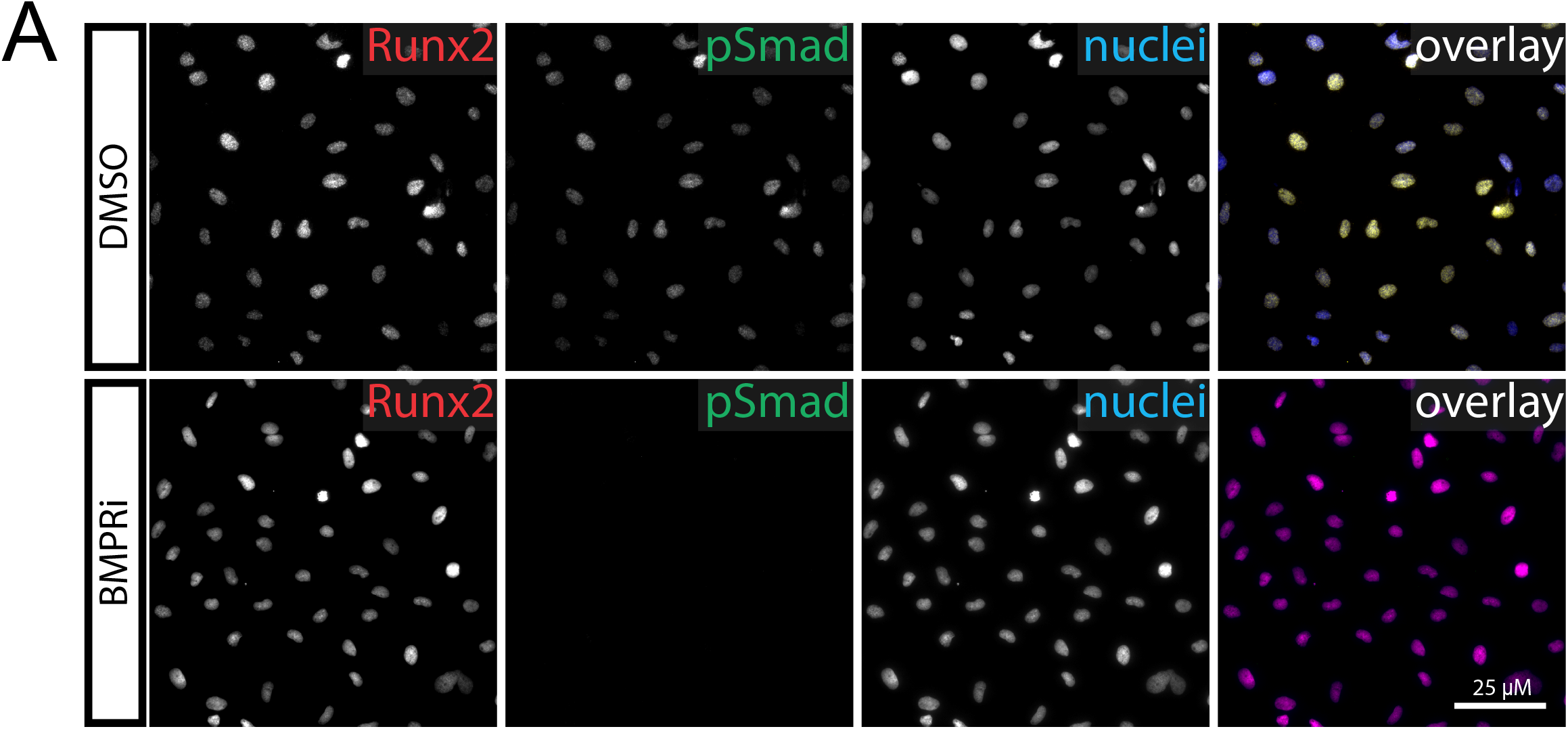
AB.9 pre-osteoblast cells intrinsically activate BMP signaling. Widefield fluorescence microscopy images of antibody-stained AB.9 cells treated with DMSO or the BMP receptor inhibitor LDN-193191. Individual channels for the osteoblast transcription factor Runx2 (red), phospho-Smad1/5/8 indicative of active BMP signaling (green), and Hoechst-stained nuclei (blue) are shown in grey scale with the overlay in color. The scale bar is 25 *μ*M.

